# Fast-annealed 3’-extended dsDNA templates facilitate efficient epitope-tag knock-in in emerging model insects

**DOI:** 10.1101/2025.06.20.660821

**Authors:** Taro Nakamura, Toshiya Ando, Yuji Matsuoka, Teruyuki Niimi

**Affiliations:** Division of Evolutionary Developmental Biology, National Institute for Basic Biology, Okazaki, Aichi, Japan, 444-8585; Basic Biology Program, The Graduate University for Advanced Studies, SOKENDAI Okazaki, Aichi, Japan, 444-8585; JST, PRESTO, Kawaguchi, Saitama 332-0012, Japan; the Hakubi Center for Advanced Research, Kyoto University, Kyoto, Kyoto, Japan, 606-8501; Graduate School of Life Sciences, Tohoku University, Sendai, Miyagi, Japan, 980-8577

**Author notes:** Corresponding authors. Correspondence and requests for materials should be addressed to T.Ni., T.A., or T.Na. These authors contributed equally to this study (Taro Nakamura, Toshiya Ando).

**Keywords:** CRISPR-Cas9, knock-in, ladybug, cricket, insect

## Abstract

CRISPR-Cas genome editing toolkits have expanded the scope of genetic studies in various emerging model organisms. However, their applications are limited mainly to knockout experiments due to technical difficulties in establishing knock-in strains, which enable *in vivo* molecular tagging-based experiments. Here, we investigated knock-in strategies in the harlequin ladybug *Harmonia axyridis*, a model insect for evolutionary developmental biology, which shows more than 200 color pattern variations within a species. We tested several knock-in strategies using synthetic DNA templates. We found that ssDNA templates generated founder knock-in strains efficiently (2.5-11%), whereas the 5’ regions of ssDNA templates were frequently deleted when the insert length exceeded ∼40 bases. To overcome this limitation, we designed several 3’ extended DNA templates. Fast-annealed 3’-extended double-stranded DNA templates, which were designed for tagging endogenous proteins with epitope tags, showed high founder generation efficiency (9.9-20.9%) and accuracy (30.8-85.7%). This strategy is also applicable to the two-spotted cricket *Gryllus bimaculatus*, suggesting that the fast-annealed 3’-extended dsDNA template is a versatile DNA template for generating knock-in strains in emerging model insects for developmental genetic studies.

**Summary statement:** Fast-annealed 3’-extended dsDNA templates facilitate efficient CRISPR-Cas9-mediated knock-in in emerging model insects.

## Introduction

The CRISPR-Cas system has enabled sophisticated genetic manipulations in model organisms, such as endogenous protein tagging (Bosch et al., 2020; Li-Kroeger et al., 2018), gene expression control (Gilbert et al., 2013; MacLeod et al., 2019; Lin et al., 2015), and cell lineage tracing (Kalhor et al., 2018). These advances have relied heavily on knock-in techniques. CRISPR-Cas approaches have also been extended to a wide range of emerging model organisms across animal and plant systems (Fei et al., 2014; Takeuchi et al., 2022; Wucherpfennig et al., 2019; Lin & Su, 2016; Chan et al., 2022; Martin et al., 2016; Zhang & Reed, 2016; Procko et al., 2023). However, in many non-model organisms, knock-in remains technically challenging due to low germline insertion efficiency and the difficulty of large-scale screening. Therefore, a versatile and efficient DNA template format that ensures high germline knock-in efficiency is urgently needed.

Traditionally, gene knock-in was achieved using linearized double-stranded DNA (dsDNA) derived from plasmids or transgenes (Thomas & Capecchi, 1987; Rong & Golic, 2000). The advent of genome-editing tools such as zinc finger nucleases, TALENs, and CRISPR-Cas systems has enabled the insertion of exogenous genes at desired sites in the genome, mainly using plasmid DNA templates (reviewed in Gaj et al. 2013). The application of genome editing tools in gene knock-in has also been achieved in various insects such as silkworm (Daimon et al., 2014; Ma et al., 2014; Zhu et al., 2015), mosquitoes (Gantz et al., 2015; Hammond et al., 2016; Kistler et al., 2015; Purusothaman et al., 2021; Rozen-Gagnon et al., 2021), and the red flour beetle (Gilles et al., 2015), although the germline insertion efficiency is often low, sometimes below 0.01 %. More recently, a single-stranded oligonucleotide (ssODN or ssDNA) template has emerged as a promising alternative, inducing more efficient knock-in in the germlines of several insects, including the fruit fly (Beumer et al., 2013), the silkworm (Ma et al., 2014; Takasu et al., 2016), and the brown bush butterfly (Connahs et al., 2022). Nevertheless, a significant drawback of the ssDNA template is the frequent occurrence of indel mutations flanking the insertion site by the double-strand break (DSB) (Bedell et al., 2012; Takasu et al., 2016; Yoshimi et al., 2016; Paix et al., 2017; Boel et al., 2018; Canaj et al., 2019; Straume et al., 2020).

Knock-in efficiency and fidelity differences stem from distinct DNA repair pathways recruited by different template types (reviewed in Yeh et al., 2019). CRISPR-induced DSBs are repaired via either precise homology-directed repair (HDR) or error-prone pathways such as non-homologous end joining (NHEJ) and microhomology-mediated end joining (MMEJ) (Rodgers & McVey, 2016). Within the HDR pathway, different sub-pathways are utilized depending on the template (Richardson et al., 2018; Gallagher et al., 2020). The dsDNA template is mainly processed by the homologous recombination repair (HRR) sub-pathway. Following a DSB, the cleaved DNA ends are resected in a 5’ to 3’ direction, creating 3’ overhangs. These overhangs are coated with RAD51 proteins, forming a filament that is crucial for invading a homologous donor template, typically the sister chromatid (reviewed in Bonilla et al., 2020). As the HRR pathway is highly active only during the S/G2 phases of the cell cycle (Karanam et al., 2012), its efficiency for integrating exogenous templates is generally low. In contrast, ssDNA templates promote DNA repair via the single-strand template repair (SSTR) pathway, a mechanism related to the single-strand annealing (SSA) sub-pathway (reviewed in Yeh et al., 2019). The SSA sub-pathway is also active during the S/G2 phases and promotes the direct annealing of complementary single-stranded DNA ends by coating them with RAD52 proteins (Liang et al., 2024). This mechanism can be exploited by exogenous ssDNA templates (reviewed in Blasiak, 2021). Therefore, donor templates need to be designed in a pathway-appropriate manner to recruit the relevant HDR subpathways efficiently. To this end, various DNA template formats, such as long ssDNA, partially annealed (melted) dsDNA, and dsDNA with a 3’-extended overhang, have been explored in cultured cells and model organisms (Yoshimi et al., 2016; Ghanta and Mello, 2020; Liang et al., 2017). However, the efficacy of these alternative templates remains largely unexplored in non-model insects.

In the present study, we aimed to identify a suitable DNA template format for non-model insects. We focused on an emerging model insect, the harlequin ladybug *Harmonia axyridis*, which exhibits over 200 color patterns within a species and is an excellent model for studying the evolutionary and developmental basis of phenotypic diversification (reviewed in Ando & Niimi, 2019). We systematically tested different DNA templates to generate knock-in strains targeting the color pattern polymorphism locus *h* (*Drosophila pannier* ortholog) in *H. axyridis*. We confirmed that ssDNA templates can efficiently generate founder knock-in individuals (2.5–11.1% efficiency), although 5’ truncations of the insert were frequently observed. Of the DNA templates tested, we found that the ‘fast-annealed’ dsDNA template with 3’ single-stranded extensions (overhang) serves as a superior template, yielding higher efficiency (9.9-20.9%) and fidelity (30.8-85.7%) in *H. axyridis*. We further observed that this strategy is effective in the two-spotted cricket *Gryllus bimaculatus*, suggesting that the fast-annealed 3’-extended DNA templates provide a useful approach for generating knock-in strains for molecular tagging-based developmental genetic experiments in emerging model insects.

## Materials and Methods

### Insect husbandry

The laboratory stock of *Harmonia axyridis* was initially established from individuals collected in Japan. They were reared at 25°C under constant light and no humidity control conditions and fed on an artificial diet (drone honeybee brood powder: sucrose: ethyl benzoate = 100: 30: 3.3) (Okada et al., 1973) or the pea aphid *Acyrthosiphon pisum*. Laboratory stock of *G. bimaculatus* was maintained in plastic cages at 26-30°C and 50% humidity under a 10 h light, 14 h dark photoperiod. They were fed on commercial fish food (Tetrafin, Spectrum Brand, USA).

### Preparation of ssDNA and CRISPR-Cas RNP complex

Synthetic CRISPR RNA (crRNA) (Table S1), trans-activating crRNA (tracrRNA), single-stranded DNA (ssDNA) templates (Table S2) with a chemical modification (Alt-R HDR modification), Cas9 protein (Alt-R® S.p. Cas9 Nuclease V3) and Cas12a protein (Alt-R® Cas12a (Cpf1) Nuclease V3) were purchased from Integrated DNA Technologies Inc. (IDT, USA). For Cas9 experiments, guide RNAs (gRNAs) were prepared by annealing crRNA and tracrRNA according to the manufacturer’s instructions. For the Cas12a experiment targeting the *G. bimaculatus piwi* locus, Cas12a crRNA was used. RNP complexes were prepared by incubating the corresponding gRNA with Cas9 or Cas12a protein at an equimolar concentration in 1× Duplex Buffer. To make Cas9 RNP, the gRNAs were prepared by annealing crRNA and tracrRNA according to the manufacturer’s instructions. Ribonucleoprotein (RNP) complexes were prepared by incubating the gRNAs and Cas9 protein at an equimolar concentration (5 µM each) in 1x Duplex Buffer (IDT, USA) at 25°C for 30 minutes. The RNP complexes were stored at -30 °C until use.

For all double-stranded DNA (dsDNA) templates, complementary oligonucleotides were annealed using a thermal cycler with the following program: 95°C for 2 min, followed by stepwise cooling to 85°C for 10 s, 75°C for 10 s, 65°C for 10 s, 55°C for 60 s, 45°C for 30 s, 35°C for 10 s, 25°C for 10 s, with a final hold at 4°C. Fast annealing was performed at a temperature ramp rate of 3.7°C/s, whereas slow annealing was performed at 0.1°C/s. Annealing was performed immediately before microinjection. The annealed dsDNA templates were mixed with pre-assembled RNP complexes immediately before injection. All templates and injection cocktails were freshly prepared for each experiment and not stored. These methods were adapted from Dokshin et al., 2018 with minor modifications.

### Generation of a long ssDNA donor template

To generate a long ssDNA donor for targeting the *h* locus, three donor fragments (i) the *h*/*pannier* left homology arm, (ii) the mCherry-P2A cassette (GenBank: AAV52164.1; Szymczak et al., 2004), and (iii) the *h*/*pannier* right homology arm were first amplified individually by polymerase chain reaction (PCR). These products were then joined using overlap-extension PCR to construct a 1.3 kb dsDNA template. This amplicon was re-amplified using a 5’-phosphorylated reverse primer to generate a substrate for strand-specific digestion, thereby rendering the antisense strand susceptible to digestion by a strand-specific nuclease. The selective digestion was performed using the Guide-it Long ssDNA Production System (Takara Bio, Japan) according to the manufacturer’s instructions. Typically, ∼5 µg of purified long ssODN was recovered from 14 µg of dsDNA template. The purified ssDNA was mixed with the RNP complex immediately before injection.

Another long ssDNA donor for targeted insertion of P2A-mNeonGreen-NLS (mNeonGreen, GenBank: AGG56535; P2A, Szymczak et al., 2004) at the C-terminus of *h*/*pannier* ORF was purchased from Integrated DNA Technologies Inc. (IDT, USA).

### Microinjection

For *H. axyridis*, microinjection was performed as previously described (Kuwayama et al., 2006; Nakamura et al., 2024). Fertilized eggs were collected within 3 hours post-oviposition for microinjection and aligned on double-sided tape affixed to a glass slide. The eggs were injected in air and kept in a moist chamber until hatching at 25°C.

For *G. bimaculatus*, microinjection was performed as previously described (Barry et al., 2019). Fertilized eggs were collected within 5 hours post-oviposition and aligned in wells of an agarose gel molded with an alignment tool. The injection mixture, containing the DNA template (1 µM) and Cas12a RNP (1 µM) in 1x Dulbecco’s Phosphate Buffered Saline (DPBS), was loaded into a glass capillary needle and injected into the fertilized eggs using a FemtoJet microinjector (Eppendorf, Germany). The eggs were injected in DPBS and kept in a moist chamber until hatching at 25°C.

### Genotyping

Genomic DNA was extracted from a batch of eggs or a single adult leg using Gencheck DNA Extraction Reagent (FASMAC, Japan) according to the manufacturer’s instructions. The target locus was amplified by PCR using KOD-FX-neo DNA polymerase (TOYOBO, Japan) and gene-specific primers (Table S3). Insertions and deletions were initially screened using a heteroduplex mobility assay on a capillary electrophoresis system with the QIAxcel Advanced System and QIAxcel DNA Screening Kit (Qiagen, Germany). The sequences of the successfully inserted alleles were determined by a Sanger sequencing service (FASMAC, Japan). Founder lines carrying insertions were initially identified by genotyping pools of 20–30 G1 eggs. The total number of screened individuals was calculated by aggregating G1 individuals derived from multiple injected egg batches. For founder lines that tested positive in the pooled-egg assay, individual G1 adults were subsequently screened by genotyping adult leg tissue.

### Estimation of G0 heritable insertion rates from sib-cross and backcross data

To estimate the proportion of G0 individuals carrying heritable insertions, we analyzed both sib-cross and backcross experiments under each condition. We defined *q* as the probability that a given G0 individual carries a heritable insertion. In backcrosses (G0 × wild type), the probability that a cross produces at least one inserted offspring is:

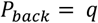

In sib-crosses (G0 × G0), a cross produces inserted offspring if at least one of the two parents is inserted:

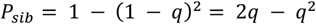

For each condition, we modeled the number of successful crosses as binomial outcomes:

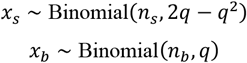

where *n*_*s*_ and *n*_*b*_ denote the number of sib-cross and backcross pairs, and *x*_*s*_ and *x*_*b*_ denote the number of crosses yielding inserted offspring.

The parameter *q* was estimated by maximum likelihood:

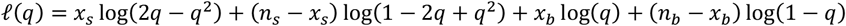

For conditions without backcross data, the terms involving *x*_*b*_ and *n*_*b*_ were omitted.

### Quantification of on-target editing by deep sequencing

To quantify on-target editing efficiency, the genomic region surrounding the CRISPR-targeted site was PCR-amplified using gene-specific primers (Table S3) and KAPA HiFi DNA Polymerase (KAPA Biosystems, USA). The amplicons were purified and size-selected using SPRIselect magnetic beads (Beckman Coulter, USA). The purified DNA libraries were subjected to paired-end sequencing on a MiSeq platform (Illumina, USA) by a commercial sequencing service (Genome-Lead, Japan). The adaptor sequences in the raw read data were trimmed using the cutadapt software (version 2.5) (Martin, 2011). The read data were mapped to the reference target sequences with and without the 3xFLAG insert and quantified using CRISPRESSO2 software (version 2.3.0) (Clement et al., 2019). The reference sequences used for read mapping and the corresponding analysis commands are provided in Data S1. To assess the average editing frequency in each experiment, we analyzed sequencing data obtained from a pooled sample of 20 fertilized eggs (N = 1). A chi-square test for pairwise comparison of indel rates between samples was performed using the pairwise.prop.test function in R (R Core team, 2021) with Holm’s correction for multiple testing (Table S4). The total reads aligned to the reference sequences were summarized in Table S6. Relative read-mapping percentages shown in Fig. 2A, B were calculated from the actual read counts provided in Table S6.

### Immunohistochemistry and image acquisition

Immunohistochemistry was performed on ladybug forewings and cricket embryos based on a previously described protocol (Patel, 1994) with some modifications. Tissues were dissected in DPBS and fixed with 4% paraformaldehyde at room temperature for 30 minutes. After washing in DPBS several times, the samples were dehydrated with methanol and stored at -30°C until use. For staining, the samples were rehydrated in DPBS, permeabilized for 30 min in DPBS containing 0.5% TritonX-100, and washed with DPBS containing 0.1% Tween 20 (PTw). After blocking for 1 hour in PTw with 1% BSA, tissues were incubated overnight at 4 °C with primary antibodies diluted in the blocking solution: rat anti-HA (Roche, Switzerland; 11867423001), mouse anti-FLAG (Sigma-Aldrich, USA; F1804, 1:200), rabbit anti-PIWI (Ewen-Campen et al., 2013, 1:300), and mouse anti-Engrailed (4D9, 1:20, Patel et al., 1989). After washing three times with PTw for 10 min each, the samples were incubated with secondary antibodies for 2 h at room temperature (Goat anti-mouse-AlexaFluor 488, Goat anti-rat-AlexaFluor 488, and Goat anti-rabbit AlexaFluor 555, Jackson ImmunoResearch Laboratories, USA; 1:200). Tissues were rewashed in PTw and mounted on glass slides with VectaShield mounting medium with DAPI (Vector Laboratories, USA). Fluorescence images were acquired using a confocal microscope (NIKON A1; NIKON, Japan). Image brightness and contrast were adjusted using the ImageJ software (Schneider et al., 2012). The final figures were assembled in Adobe Illustrator software (Adobe, USA).

**Table 1.**
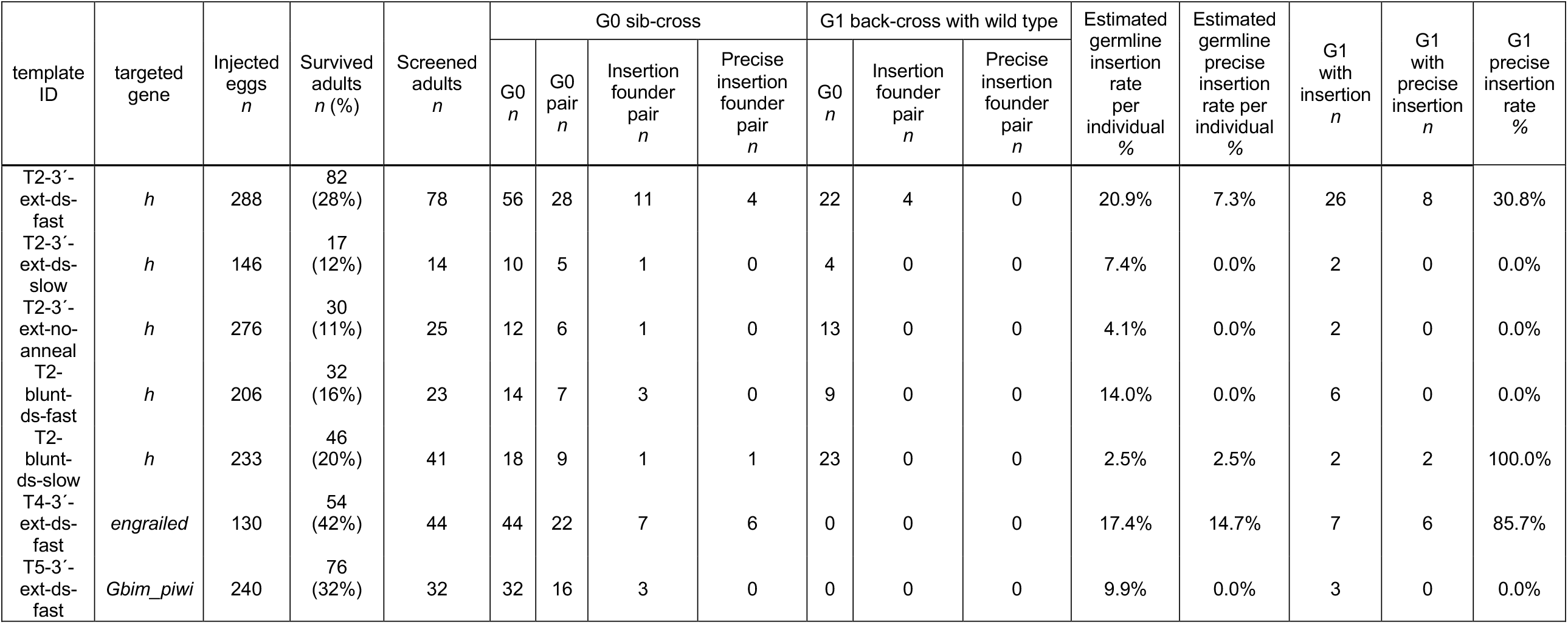
Statistics of the gene knock-in using dsDNA templates.

## Results and Discussion

### Targeted gene disruption by the CRISPR/Cas9 system in *H. axyridis*

To establish genome editing in *H. axyridis*, we injected Cas9 protein and single guide RNA (sgRNA) targeting the elytral color-patterning gene *h* (*Drosophila pannier* ortholog) into fertilized eggs (Gautier et al., 2018; Ando et al., 2018). Although CRISPR-mediated knockout has been reported previously (Wu et al., 2022; Partosh et al., 2024), we established our protocol based on our laboratory’s transgenesis and TALEN methods (Kuwayama et al., 2006; Hatakeyama et al., 2016). sgRNA targeting the 5′ end of the *h* ORF was designed in the dominant *h*^*C*^ allele (Fig. 1A, T1) to induce frameshift mutations. Of 108 injected eggs, 18 larvae hatched, and nine showed mosaic elytral color loss (Fig. 1B, G0 crispant). Mutant strains were established by crossing G0 adults with each other or with wild type. Homozygous *h*^*C*^ mutants were lethal, and complementation tests with the recessive *h* allele confirmed the loss-of-black-pigment phenotype consistent with *h*/*pannier* RNAi mutants (Fig. 1B, *h*^*C-del*^/*h*). These results confirmed successful establishment of CRISPR/Cas9 genome editing in *H. axyridis*.

**Figure 1.**
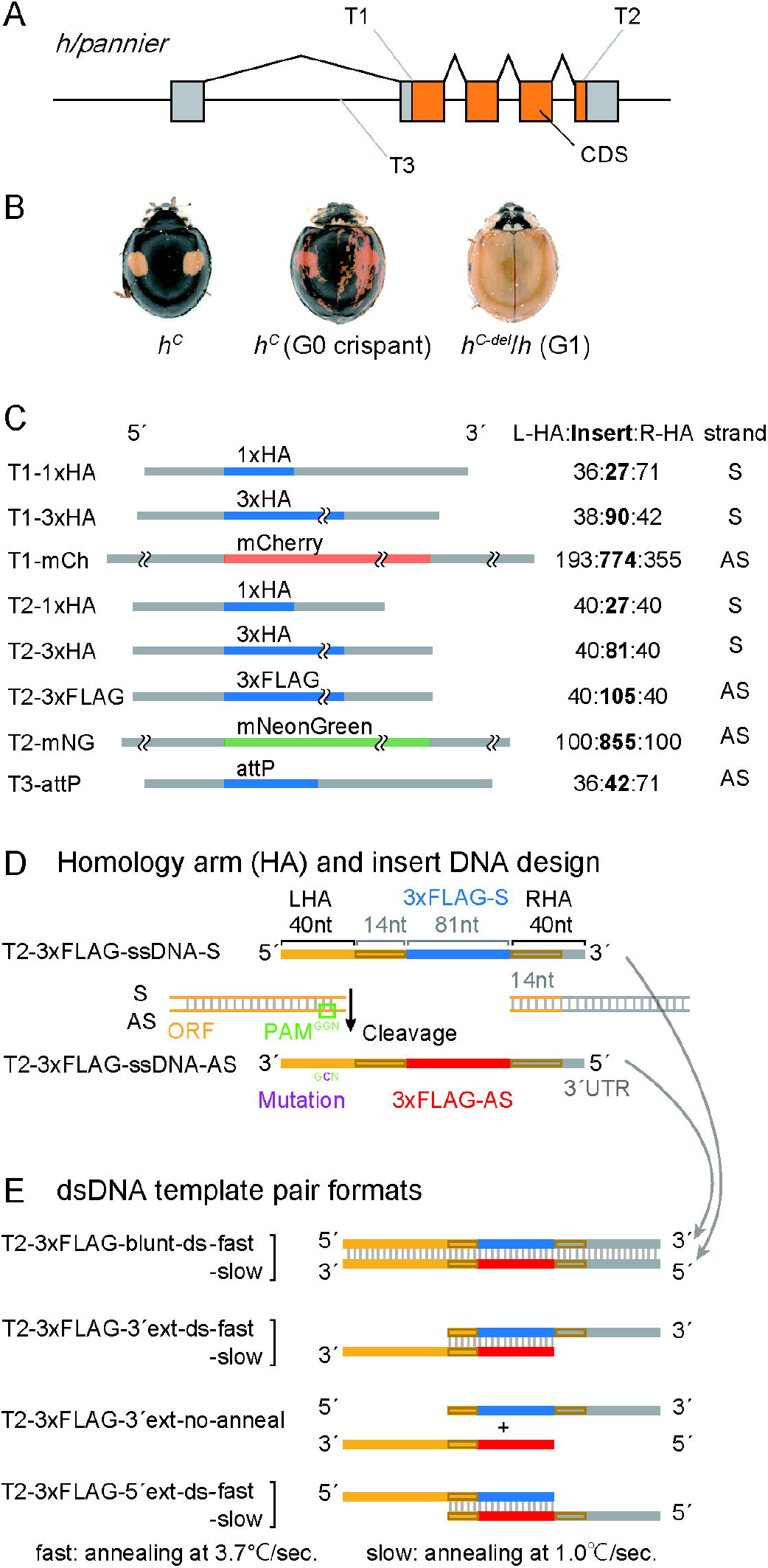
The experimental design of the gene knockout and the ssDNA-mediated gene knock-in targeting the *h*/*pannier* gene in *H. axyridis*. (A) The exon-intron structure of the *h*/*pannier* gene and the CRISPR/Cas9 target sites. Target 1 (T1): the 5’ end of the coding sequence [CDS], Target 2 (T2): the 3’ end of CDS, Target 3 (T3): one of the breakpoints of the ancestral inversion within the first intron [Ando et al., 2018]). (B) The representative phenotypes of *h* knockout individuals. *h*^*C*^: wild-type allele showing two red/orange spots on a black background. G0 crispants: a patchy reduction of black pigmentation was observed. *h*^*C-del*^ /*h* (G1): heterozygous mutants carrying the recessive *h* allele lacked black spots. (C) The schematics of the ssDNA templates that we used in the knock-in experiments. The 5’ ends are on the left side, and the 3’ ends are on the right. The length of the left homology arm (LHA), the insert (Insert), the right homology arm (RHA), and the strand of ssDNA against the gene (S: sense and AS: antisense) are indicated on the right. (D) Base design of sense- and antisense-strand ssDNA knock-in templates (T2-3xFLAG-ssDNA-S and T2-3xFLAG-ssDNA-AS). The templates consisted of 40 nt left and right homology arms (LHA and RHA) corresponding to the cleaved terminal sequences, together with an insert sequence containing the deleted 14 nt ORF fragment, GS linker, and 3×FLAG tag sequences. (E) dsDNA templates used in the experiments. T2-3xFLAG-blunt-ds: blunt-ended dsDNA template. T2-3xFLAG-3′ext-ds: 3′-extended dsDNA template. T2-3xFLAG-5′ext-ds: 5′-extended dsDNA template. “-fast” and “-slow” indicate annealing rates of 3.7 °C/s and 0.1 °C/s, respectively. Sense and antisense strands are shown in blue and red, respectively.

**Figure 2.**
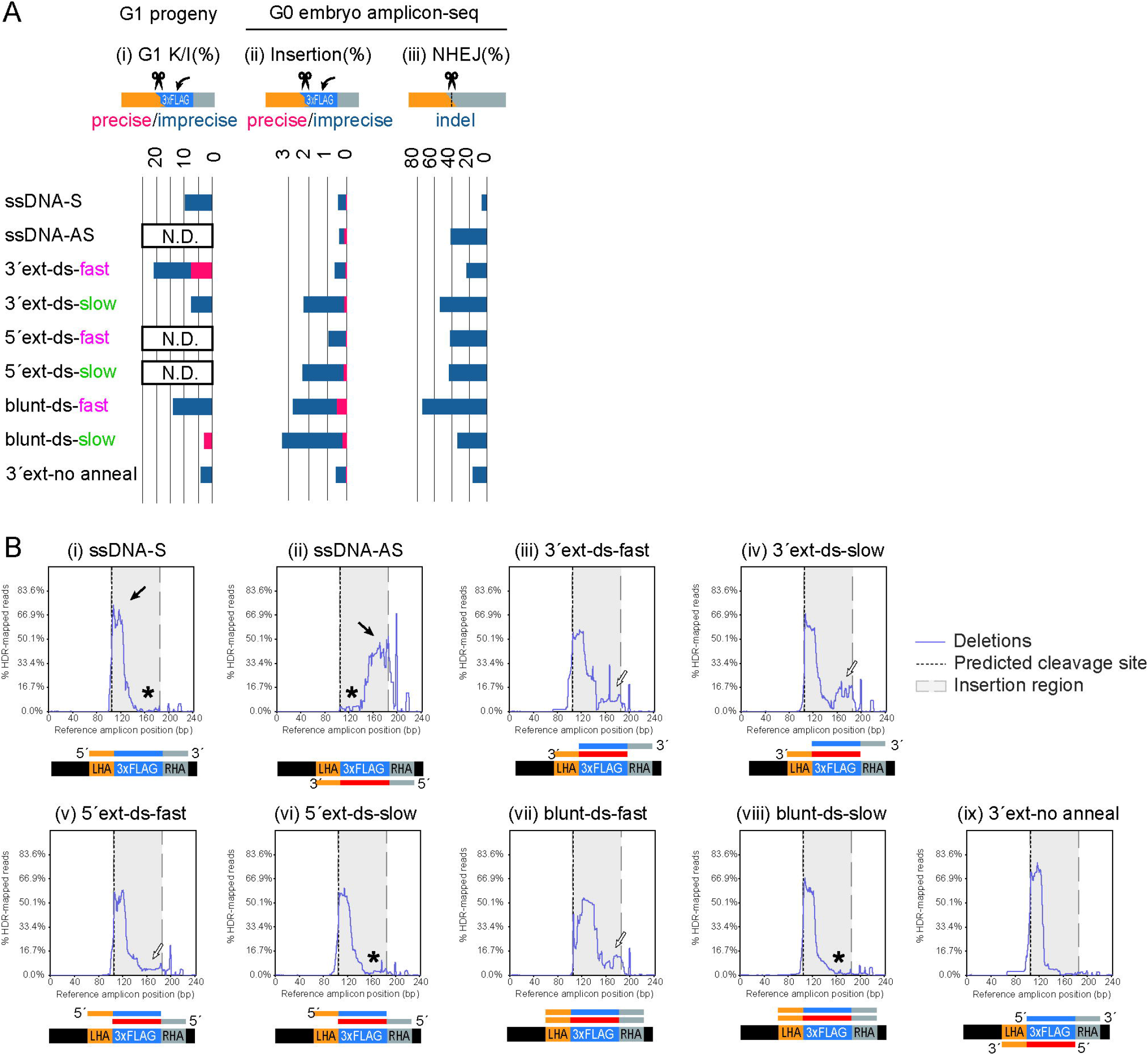
Indel patterns in knock-in-treated *H. axyridis* embryos and their offspring. (A) Insertion and indel rates in the G1 progeny and G0 embryos injected with different ssDNA or dsDNA templates. The template names correspond to those in Fig. 1D and Fig. 1E, with the “T2-3xFLAG” prefix omitted. Bar graph (i), G1 K/I (Knock-In): percentage of the G1 progenies (bar graph showing the estimated germline precise insertion rate per individual (%) from Table 1). Red: precise insertions. Blue: imprecise insertions. N.D., not determined. Bar graph (ii), insertion frequencies of precise (red) and imprecise (blue) 3xFLAG insertions in amplicon-seq data from G0 embryos. Bar graph (iii), NHEJ without knock-in insert: percentage of indel reads lacking the 3xFLAG insert in G0 embryo amplicon-seq data. (B) Deletion patterns of 3xFLAG-inserted reads in the deep sequencing analysis of the G0 embryos. x-axis: reference sequence position; y-axis: deletion frequency at each site; actual read counts are provided in Table S6. Black arrows, deletions corresponding to the 5’-region of ssDNA templates (Mean deletion frequency >33.3% within 25-nt insert termini). White arrows, a slight increase in deletions near the 3’ end of inserts (Mean deletion frequency 3.84-27.0% within 25-nt insert termini). Asterisk: low frequency of deletions near the 3’ end of inserts in the slowly annealed templates (Mean deletion frequency <1.2% within 25-nt insert termini). The schematics of the reference sequence (LHA, 3xFLAG, and RHA) and the injected DNA template format are depicted in all charts in this figure. Shaded area, 3xFLAG insert region; dashed line, predicted Cas9-RNP cleavage site.

### ssDNA-mediated precise insertion in *H. axyridis*

Efficient ssDNA-mediated knock-in methods have recently been reported in multiple organisms (Chen et al., 2011; Yoshimi et al., 2016; Quadros et al., 2017; Kanca et al., 2019; Inoue et al., 2021; Heryanto et al., 2022; Connahs et al., 2022). We tested chemically modified ssDNA templates carrying small tags (1×HA, 3×HA, 3×FLAG) or an attP site at three positions in the *h* gene: the 5′ end of the ORF (T1), the 3′ end of the ORF (T2), and the first intron (T3) (Fig. 1A) (Fig. 1C, T1-1xHA, T1-3xHA, T2-1xHA, T2-3xHA and T2-3xFLAG, and T3-attP). Left/right homology arm lengths were 36/71 nt, 38/42 nt, 118/40 nt, or 40/40 nt; some templates used asymmetric homology arms, which were reported to enhance insertion efficiency in human cells (Richardson et al., 2016). Cas9 RNP (5 µM) and ssDNA templates (5 µM) targeting the antisense strand of *h* were injected into embryos. Germline-transmitted insertions were obtained in most cases (Table S5, Insertion founder pair, 2.5% [1/40]–11% [1/9]), except for T2-1xHA (0% [0/14]). However, precise insertions were detected only in two strains (Table S5, Precise insertion, T1-1xHA and T3-attP). Longer inserts, including mCherry and P2A-mNeonGreen-NLS, also failed to yield precise insertions (Fig. 1C; Table S5, Precise insertion, T1-mCh and T2-mNG). These results suggested that precise insertion occurred only with inserts ≤42 nt (Fig. 1C; Table S5, “Precise insertion”, T1-HA and T3-attP). Importantly, even in imprecise insertion strains, the 3′ ends of ssDNA templates were inserted with high fidelity (Table S5, 3′ Fidelity, 80% [4/5]–100% [1/1–5/5], except for T1-mCh and T2-mNG [0/1]), whereas the 5′ ends were frequently truncated (Table S5, 5′ Fidelity, 0% [0/1–0/5]–80% [4/5]). Based on this finding, we designed templates flanked by high-fidelity 3′ ends on both sides of the insert.

### Fast-annealed 3’-extended dsDNA template confers efficient and accurate knock-in outcomes

To improve insertion fidelity at both ends of the insert, we designed dsDNA templates with 3′-extended homology arms targeting the same CRISPR/Cas9 cleavage site at the 3′ end of the *h*/*pannier* ORF (Fig. 1A,D,E; Fig. S1). As the insert, we used a 3×FLAG tag (81 nt including a GS-linker). Donor templates contained 40-nt homology arms flanking the cleavage site, a truncated 14-nt ORF fragment, and a mutated PAM sequence to prevent re-cleavage after insertion (Fig. 1D). We compared 3′-extended, blunt-ended, and no-annealed templates prepared under fast- or slow-annealing conditions (Fig. 1E; fast-annealing: 3.7 °C/s; slow-annealing: 0.1 °C/s). Because fast annealing reportedly enhances knock-in efficiency in *Caenorhabditis elegans* (Ghanta and Mello, 2020), both annealing conditions were tested. All dsDNA templates were generated by annealing ssDNA oligonucleotides.

Among them, the fast-annealed 3′-extended dsDNA templates showed the highest insertion efficiency (∼20.9% [11/78]) and precise insertion rate (30.8% [8/26]) in G1 offspring (Table 1, T2-3xFLAG-3’ext-ds-fast; Fig. 2A). Moreover, fast-annealed templates yielded more efficient insertions than slowly annealed or no-annealed templates (Table 1, Estimated germline insertion rate per individual, T2-3′-ext-ds-fast, ∼20.9% [11/78]; T2-blunt-ds-fast, ∼14.0% [3/23] versus T2-3′-ext-ds-slow, ∼7.4% [1/14]; T2-blunt-ds-slow, ∼2.5% [1/41]; T2-3′-ext-no-anneal, ∼4.1% [1/25]) (Fig. 2A(i)). Therefore, a fast-annealed format would be a better design for efficient insertion in the germline.

On the other hand, with respect to insertion precision, slowly annealed blunt-ended dsDNA yielded one precise insertion strain (∼2.5% [1/41]), although its overall insertion efficiency was low. In contrast, fast-annealed blunt-ended dsDNA failed to produce precise insertions (0% [0/23]) despite relatively efficient imprecise insertion (∼14% [3/23]), suggesting that the combination of blunt-ended templates and fast annealing does not favor efficient and precise insertion. Therefore, fast-annealed 3′-extended dsDNA templates provide a more suitable template design for efficient and precise germline insertion. All insertion patterns identified in offspring are summarized in Fig. S2.

### DNA repair patterns associated with highly efficient and precise knock-in

To gain insights into the molecular features associated with efficient and precise insertion of fast-annealed 3′-extended dsDNA, we deep-sequenced the target region at the 3′ end of the *h* ORF (Fig. 1D, T2) in G0 embryos injected with various knock-in templates, including ssDNA templates (sense, antisense, or unannealed mixed strands for 3′-extended dsDNA) and dsDNA templates (3′-extended, 5′-extended, or blunt-ended) prepared under fast- or slow-annealing conditions (Fig. 1E). Amplicon-seq reads from pooled injected embryos (20 embryos; N = 1 per template) were mapped to reference sequences with or without the insert, and indel patterns were quantified (Fig. 2A,B; Table S6).

In G0 embryos, fast-annealed blunt-ended dsDNA templates showed the highest precise insertion frequency (0.52%, 381/73,169 reads), whereas fast-annealed 3′-extended dsDNA templates showed only 0.01% (9/64,634 reads) (Fig. 2A(ii); Tables S4 and S6). Even when imprecise insertions were included, 3′-ext-fast was not the most efficient condition in G0 embryos (Fig. 2A(ii), blue + red bars). These results indicate that repair outcomes in G0 embryos do not necessarily reflect knock-in efficiency in G1 offspring, potentially because G0 embryos mainly reflect somatic repair, whereas G1 offspring reflect germline repair, including germline-specific DNA damage responses such as p53-mediated apoptosis (Gartner et al., 2000). Accordingly, G0 insertion efficiency alone may not reliably predict knock-in efficiency in the G1 generation.

Nevertheless, some insertion patterns in G0 embryos were consistent with those observed in G1 offspring. Notably, in ssDNA-injected embryos, the 3′ region of the 3×FLAG insert was frequently retained (Fig. 2B(i,ii), asterisks), whereas the 5′ region was often deleted, mirroring the insertion patterns observed in G1 progeny (Fig. 2B(i,ii), black arrows). In addition, fast-annealed templates associated with efficient G1 insertion (Table 1, T2-3′-ext-ds-fast and T2-blunt-ds-fast) showed moderately elevated deletion frequencies within the inserted 3×FLAG tag (Fig. 2B(iii,iv,vii), white arrows), whereas slow-annealed templates showed near-complete insertions with minimal deletions (Fig. 2B(vi,viii), asterisks). These deletions may result from mismatch repair triggered by partially annealed inserts (Li, 2008), suggesting that moderate deletion frequencies in G0 embryos reflect repair processes facilitating efficient germline insertion. This pattern might reflect elevated background activity of error-prone SSA-mediated repair in embryos, increasing sporadic precise insertion events in germline-contributing cells.

Furthermore, indel-containing reads lacking the 3×FLAG insert, indicative of NHEJ-mediated repair, were less frequent with fast-annealed 3′-extended dsDNA (22.5%) than with other annealed dsDNA templates (33.9–73.9%; Fig. 2A(iii); Table S4; chi-square test with Holm’s correction). NHEJ frequencies were further reduced in ssDNA-S-injected embryos (5.4%), which also showed efficient G1 insertion (9.8%; Fig. 2A(i,iii); Tables S5 and S6). These results suggest that 3′-extended homology arms facilitate annealing to resected DNA ends and suppress NHEJ-mediated repair. Consistently, blunt-ended templates showed high NHEJ frequencies (74% in blunt-fast and 34% in blunt-slow; Fig. 2A(iii)). Thus, reduced NHEJ signatures at the cleavage site may reflect repair conditions favorable for efficient G1 insertion.

Taken together, the combination of moderate insert deletions and reduced NHEJ signatures in G0 sequencing data may predict efficient and precise G1 insertion conditions. This strategy may reduce the need for extensive injection screening in newly studied insect species.

### Fast-annealed 3’-extended dsDNA templates as a universal template for efficient knock-in insects

We further confirmed the utility of the fast-annealed 3’-extended dsDNA format by targeting another gene, the wing posterior compartment determinant gene *engrailed*. With this approach we observed highly efficient (∼17.4% = ∼7/44) and precise (85.7% = 6/7) insertion of the 3xFLAG tag (Table 1, T4-3’-ext-ds-fast; Fig. S3). We confirmed the utility of the knock-in tags by immunohistochemistry against the inserted tags. We successfully detected the expected expression pattern of the target genes in developmental tissues—H/Pannier in presumptive black spots in the pupal wing, where red pigment granules do not accumulate (Fig. 3A) (Ando et al., 2018; Gautier et al, 2018), and Engrailed in larval wing discs in the presumptive posterior region (Fig. 3B)—thus demonstrating the feasibility of this method for molecular developmental studies.

**Figure 3.**
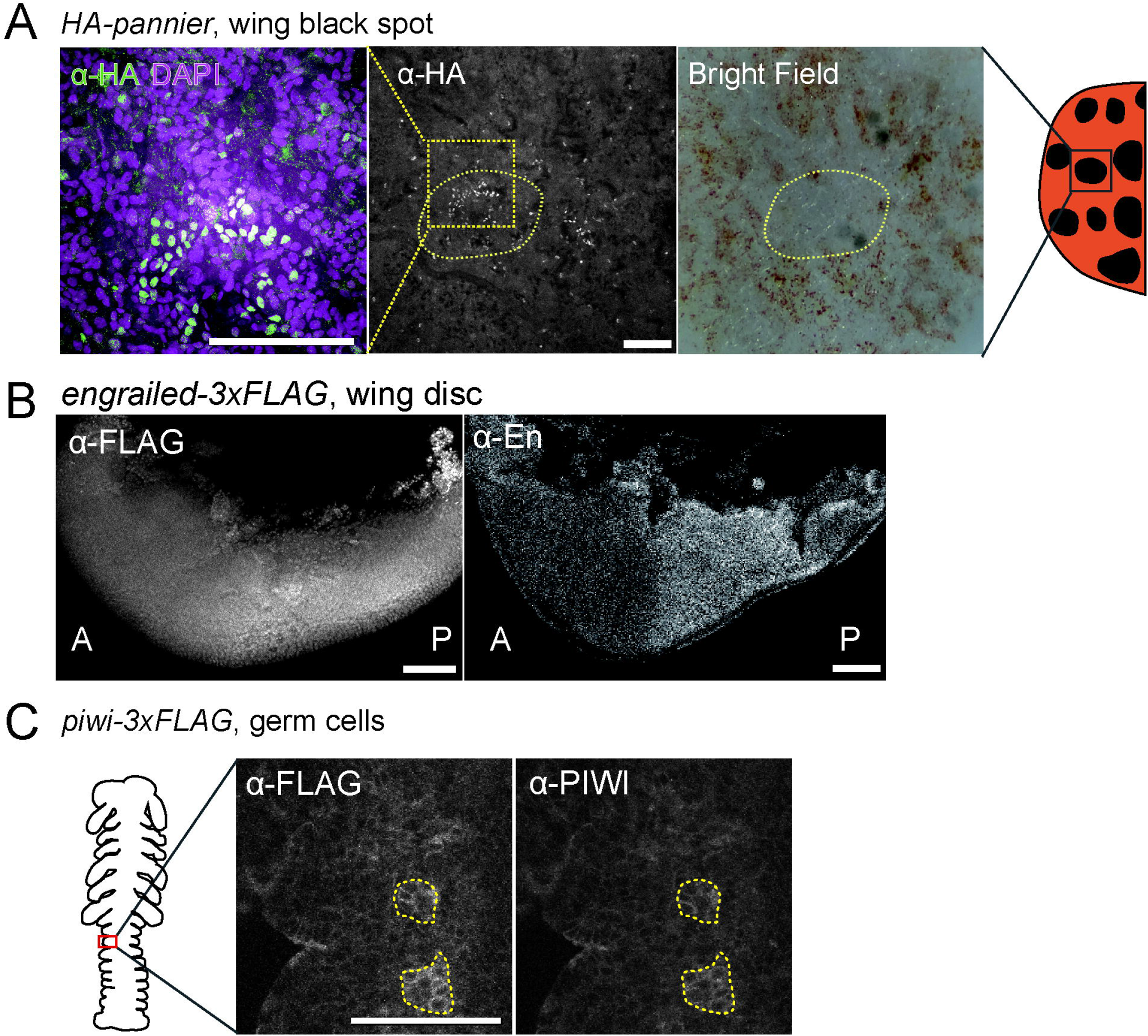
Immunofluorescence detection of endogenous protein using knock-in epitope tags. (A) H/Pannier protein expression in a presumptive black pigment region of the *H. axyridis* pupal forewing. Left, merged fluorescence image; middle, α-HA channel; right, bright-field image. Yellow dashed outlines indicate a presumptive black pigment cell cluster lacking red pigment granules. Green, α-HA; magenta, DAPI. (B) Engrailed (En) protein expression in a *H. axyridis* larval wing disc. Left, α-FLAG staining in the *engrailed-3xFLAG* knock-in line; right, α-En staining. A, the presumptive anterior compartment. P, the presumptive posterior compartment. (C) Piwi protein expression in primordial germ cells of a *G. bimaculatus* embryo. Piwi–3xFLAG was detected using anti-FLAG antibody and compared with anti-Piwi staining. Yellow dashed outlines indicate representative Piwi-positive primordial germ cells. Scale bars, 100 µm (A-C).

To test cross-species utility, we applied the same approach to the two-spotted cricket *Gryllus bimaculatus*, which diverged from the common ancestor with the ladybug ∼390 Ma (Misof et al., 2014). We designed a targeted 3xFLAG tag insertion at the 3’ end of the germline-specific gene, *piwi* (Ewen-Campen et al., 2013), using Cas12a/Cpf1 (Zetsche et al., 2015) due to PAM constraints near the targeted site at the *piwi* locus. Efficient insertion of 3xFLAG tag was also achieved (∼9.9% = ∼3/32; Table 1, T5-3’-ext-ds-fast). However, in all obtained strains (n = 3; Table 1, T5-3’-ext-ds-fast), the 3’ end of the endogenous target sequence was duplicated (Fig. S4A, red bars), likely because the 5’ overhangs generated by Cas12a cleavage were not degraded during repair. Despite the sequence duplication, the epitope tag remained in frame because the duplicated fragment was outside the CDS. Moreover, homozygotes were viable, suggesting that the inserted tag encoded a functional PIWI protein, as detected by FLAG immunostaining (Fig. 3C; Fig. S3B, yellow broken lines). These results demonstrate that this strategy can be applied to diverse insect species for tagging endogenous proteins.

## Conclusions

We demonstrated that the fast-annealed 3’-extended dsDNA templates are functional knock-in templates that can induce precise and efficient insertion at the targeted site and overcome the frequent insert deletion observed in *H. axyridis*. This template design can be also applied to a distantly related insect, the two-spotted cricket *Gryllus bimaculatus*, suggesting that this DNA template format would be a versatile knock-in template for insects. The future challenge to be tested is whether much longer DNA, such as that coding the *GFP* gene, can be inserted into the targeted site using our DNA template. To this end, the cost of DNA synthesis must be much lower to inject fertilized eggs enough to obtain progenies with inserts. Our strategy will be a practical approach to accessing sophisticated CRISPR-mediated developmental genetic tools in emerging model insects.

## Author contributions

T.Na, T.A, and T.Ni. conceived the project. T.Na, T.A., and Y.M. designed DNA templates for knock-in. T.Na and T.A performed micro-injection experiments in *H. axyridis*. T.Na and Y.M performed immunohistochemical analysis in *H. axyridis*. T.Na performed all experiments in *Gryllus bimaculatus*. T.A. performed data analysis for deep sequencing. T.A. wrote the paper. All the authors commented on the manuscript. T.A. finalized the manuscript with input from other authors.

## Acknowledgments

We thank Ms. Junko Morita for experimental support in DNA sample preparation and genotyping, Ms. Haruka Kawaguchi, Ms. Rie Taguchi, Ms. Michiko Yokoyama, and Ms. Hiroko Sugiyama for insect husbandry. We thank the Emerging Model Organisms Facility of the NIBB Trans-Scale Biology Center for technical assistance. The anti-Piwi antibody was a gift from Dr. Cassandra Extavour. The anti-Engrailed antibody developed by Patel et al. (1989) was provided by the Developmental Studies Hybridoma Bank, created by the NICHD of the NIH and maintained at The University of Iowa, Department of Biology, Iowa City, IA 52242. This work was supported by the NIBB Collaborative research projects for integrative genomics (23NIBB402). English editing was supported by Grammarly and ChatGPT-5.

## Data availability

The raw read data for deep sequencing analysis was deposited at GenBank with the accession number PRJNA1257362.

