## Supplementary Information for "Fast-annealed 3’-extended dsDNA templates facilitate efficient epitope-tag knock-in in emerging model insects"

### Supplementary Materials and Methods

#### Supplementary Data

**Data. S1**

amplicon_with_no_insert="AACCAAGGATACCCTTACCAGCAAATCTTCGGGTTCCCAGGTG
CTGGACCGACAAACCCAGAATTAGCGTATCATCACCAACATCACGTAACAGCTTCTGCCA
AACTGATGGCTACGACATAAACAAGACAAGACAAGACTGGGAGAATCAACAAAGACAC
T"
amplicon_with_expected_insertion="AACCAAGGATACCCTTACCAGCAAATCTTCGGGTTCCC
AGGTGCTGGACCGACAAACCCAGAATTAGCGTATCATCACCAACATCACGTAACAGCTTC
TGCAAAACTGATGGCTACGACAGGAGGGTCCGGAGGCGACTACAAAGACCATGACGGT
GATTATAAAGATCATGATATCGATTACAAGGATGACGATGACAAGTAAACAAGACAAGAC
AAGACTGGGAGAATCAACAAAGACACT"

```
CRISPResso -r1 "$read_file_1" -r2 "$read_file_2" ¥
   -a "$amplicon" ¥
   -e "$expected" ¥
   -g TTATGTCGTAGCCATCAGTT ¥
   --plot_histogram_outliers
```

**Supplementary Figures**

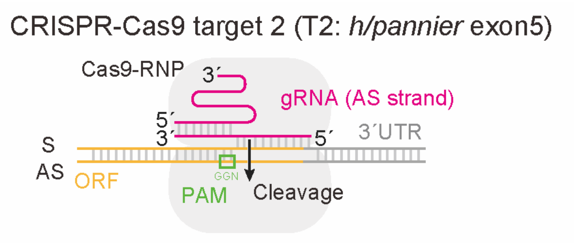


**Fig. S1 Schematics of the T2 cleavage site used to evaluate knock-in efficiency across different DNA template formats.** Orange regions indicate the ORF of the *h/pannier* gene, whereas gray regions indicate the 3′-UTR of *h/pannier*. Green indicates the PAM sequence, and magenta indicates the gRNA. The CRISPR/Cas9 protospacer sequence and the protospacer adjacent motif (PAM) were designed on the antisense strand at the 3´ end of the *h/pannier* ORF.

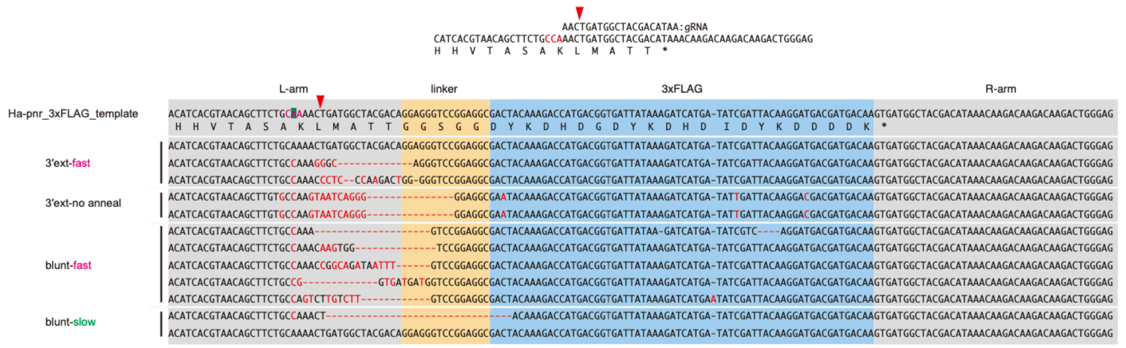


**Fig. S2. Sequence analysis of CRISPR-mediated 3xFLAG knock-in alleles at the *Ha-h/pannier* locus.** Alignment of the wild-type target sequence, donor template, and representative knock-in alleles obtained by CRISPR-mediated insertion of a linker–3xFLAG cassette at the *Ha-h/pannier* locus. The gRNA target sequence and the predicted Cas9 cleavage site are shown at the top, with the red arrowhead indicating the predicted double-stranded DNA breakpoint. The donor template (Ha-pnr_3xFLAG_template) consists of the left homology arm, linker sequence, 3xFLAG coding sequence, and right homology arm, indicated by gray, orange, blue, and gray shading, respectively. Knock-in alleles recovered under the indicated template-preparation conditions are shown below the donor template. Nucleotides matching the donor template are shown in black, whereas substitutions, insertions, or other sequence differences are highlighted in red. Dashed lines indicate gaps introduced in the alignment, corresponding to deletions or missing sequence relative to the donor template.

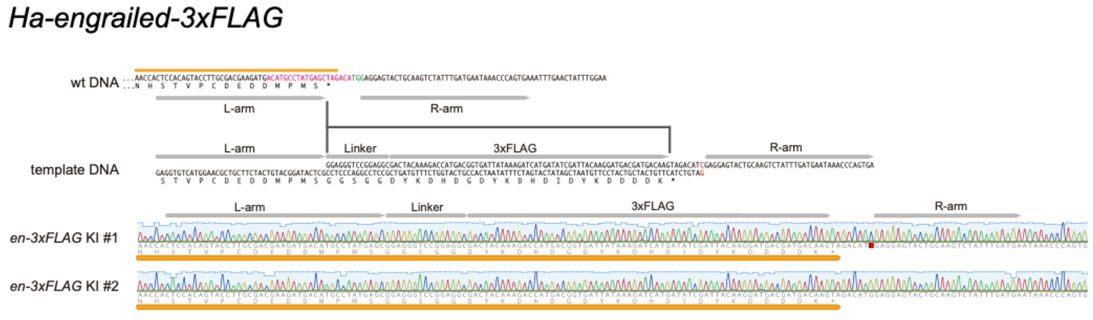


**Fig. S3. CRISPR-mediated knock-in of a 3xFLAG tag at the *Ha-engrailed* locus.**

Schematic representation of the CRISPR/Cas9-mediated C-terminal 3xFLAG knock-in strategy for *Ha-engrailed* (*Ha-en*). The wild-type genomic sequence around the end of the coding region of *Ha-engrailed* is shown at the top, with the PAM indicated in green and the sgRNA target sequence in magenta. The donor template contains left and right homology arms flanking an in-frame linker–3xFLAG cassette, which was inserted immediately upstream of the translation termination site. The original PAM site was mutated to avoid unwanted recutting of the genome and donor sequences, as indicated in red. The resulting allele encodes Ha-Engrailed fused to a C-terminal GGS linker and 3xFLAG tag. Sanger sequencing chromatograms of two independent knock-in samples, *en-3xFLAG* KI #1 and #2, confirm precise insertion of the linker–3xFLAG sequence at the intended genomic position. The *en-3xFLAG* KI #1 allele carries a mutated PAM, whereas KI #2 retains the original PAM. Subsequent experiments were performed using en-3xFLAG KI #1.

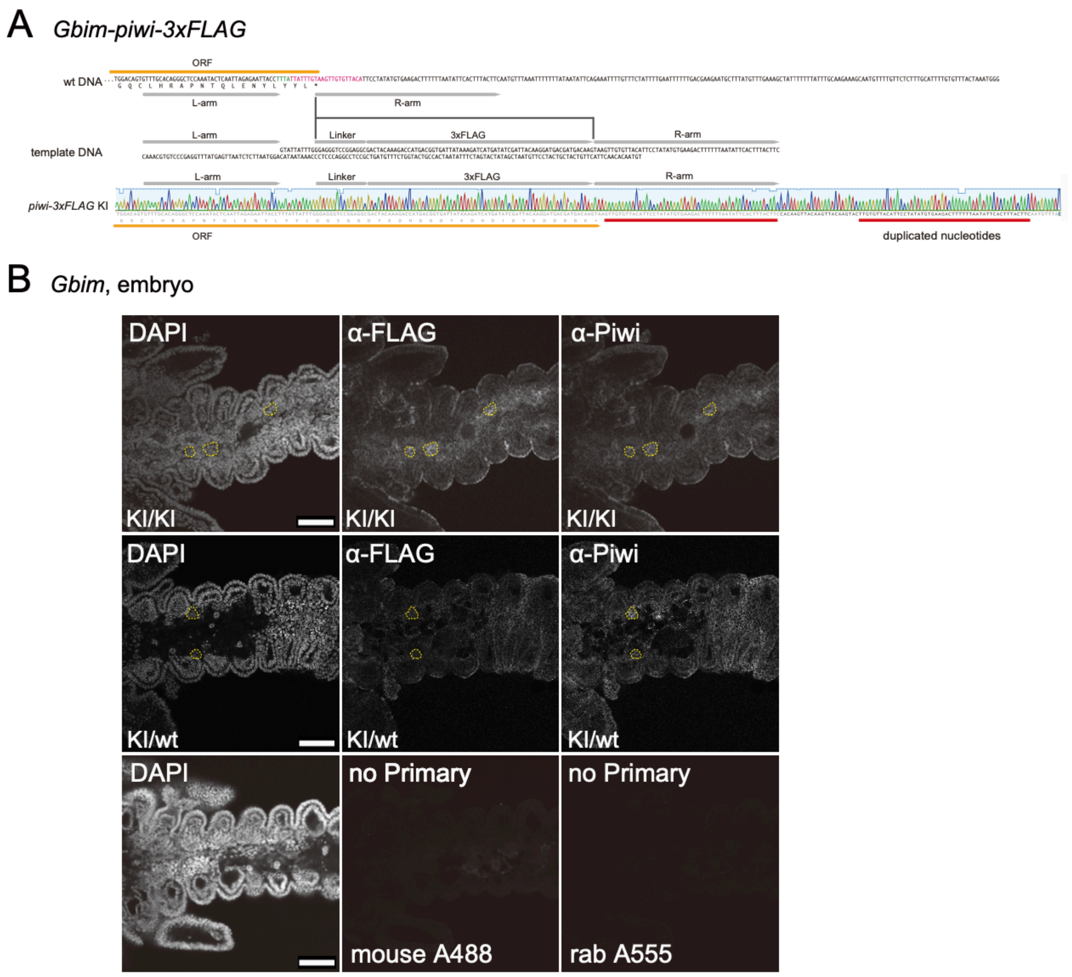


**Fig. S4. Generation and validation of the *Gbim-piwi–3xFLAG* knock-in line.**

(A) Schematic representation of the CRISPR-mediated 3xFLAG tag knock-in strategy at the *G. bimaculatus piwi* locus. The wild-type genomic sequence, donor template containing the left and right homology arms, linker, and 3xFLAG tag, and the resulting *piwi–3xFLAG* knock-in allele are shown. The target/insertion site is indicated, and Sanger sequencing of the knock-in allele confirms insertion of the linker–3xFLAG sequence. Duplicated nucleotides detected adjacent to the insertion are indicated in the UTR region.

(B) Immunohistochemical validation of Piwi–3xFLAG expression in *G. bimaculatus* embryos. Nuclei were stained with DAPI, and Piwi–3xFLAG was detected using anti-FLAG antibody together with anti-Piwi antibody. The upper images show the uncropped fields corresponding to the cropped images presented in the main figure. The upper row shows homozygous knock-in embryos (KI/KI), and the middle row shows heterozygous knock-in embryos (KI/wt). The lower row shows no-primary-antibody negative controls imaged in the mouse Alexa Fluor 488 and rabbit Alexa Fluor 555 channels. No specific signal was detected in either channel. Yellow dashed circles indicate representative Piwi-positive cells, which are primordial germ cells. Scale bars, 100 μm.

#### Supplementary Tables

Table S1. gRNA target sequences used in this study

| Name | gRNA target sequence | targeted gene | target region | purpose |
|---|---|---|---|---|
| Haxy_h_5p | GGGACTGCTGAAGGTGTTGG(TGG) | *h* | 5' | KO/KI |
| Haxy_h_3p | TTATGTCGTAGCCATCAGTT(TGG) | *h* | 3' | KI |
| Haxy_h_intron | AGCAATCGTGCAGGTGCAGT(TGG) | *h* | intron | KI |
| Haxy_en | ACATGCCTATGAGCTAGACA(TGG) | *engrailed* | 3' | KI |
| Gbim_piwi_3p_Cas12a | (TTTA)TTATTTGTAAGTTGTGTTACA | *piwi* | 3' | KI |

### Table S2. DNA templates used for targeted gene knock-ins

| ID | template ID | Species | targeted gene | sequence | insert tag | ssDNA strand | ssDNA length | LHA length (nt) | Insert length (nt) | RHA length (nt) | distance from cleavage end to inner LHA end | distance from cleavage end to | gRNA strand |
|---|---|---|---|---|---|---|---|---|---|---|---|---|---|
| Haxy-h-5-1xHA | T1-1xHA | *Harmonia axyridis* | *h* | AACGGCTATGGCGACGGCAACGCGGGTTTCCACCAATACCCATATGATGTTC<br>CTGACTATGCGCACCTTCAGCAGTCCCCCGTCTACGTGCCGAGCAGCAGGG<br>CCGTACCGCACCAATATTCCCCGGCGGCAGG | 1xHA | S | 134 | 36 | 27 | 71 | 1 | -2 | AS |
| Haxy-h-5-3xHA | T1-3xHA | *Harmonia axyridis* | *h* | GCAACGGCTATGGCGACGGCAACGCGGGTTTCCACCAATACCCATACGATG<br>TTCCTGACTATGCGGGCTATCCCTATGACGTCCCGGACTATGCAGGATCCTA<br>TCCATATGACGTTCCAGATTACGCTCACCTTCAGCAGTCCCCCGTCTACGTG<br>CCGAGCAGCAGGGCC | 3xHA | S | 170 | 38 | 90 | 42 | 1 | -2 | AS |
| mCherry-2A | T1-mCh | *Harmonia axyridis* | *h* | AGTTATTCTTGAACGGTCACCCATCCAGATATTGACCACGTCTAAAGCTTCTT<br>AGCTTCCAAATATTTAAGCCGAATTTCAATAATTCCATTTTTCTTTTTTCCAGG<br>TCCGTAATTAGCCCGAACAGGCCAGCATGTTCCACACCGGCGCCGGCGGCA<br>ACGGCTATGGCGACGGCAACGCGGGTTTCCACCAAATGGTGAGCAAGGGCG<br>AGGAGGATAACATGGCCATCATCAAGGAGTTCATGCGCTTCAAGGTGCACAT<br>GGAGGGCTCCGTGAACGGCCACGAGTTCGAGATCGAGGGCGAGGGCGAGG<br>GCCGCCCCTACGAGGGCACCCAGACCGCCAAGCTGAAGGTGACCAAGGGT<br>GGCCCCCTGCCCTTCGCCTGGGACATCCTGTCCCCTCAGTTCATGTACGGC<br>TCCAAGGCCTACGTGAAGCACCCCGCCGACATCCCCGACTACTTGAAGCTGT<br>CCTTCCCCGAGGGCTTCAAGTGGGAGCGCGTGATGAACTTCGAGGACGGCG<br>GCGTGGTGACCGTGACCCAGGACTCCTCCCTGCAGGACGGCGAGTTCATCT<br>ACAAGGTGAAGCTGCGCGGCACCAACTTCCCCTCCGACGGCCCCGTAATGC<br>AGAAGAAGACCATGGGCTGGGAGGCCTCCTCCGAGCGGATGTACCCCGAG<br>GACGGCGCCCTGAAGGGCGAGATCAAGCAGAGGCTGAAGCTGAAGGACGG<br>CGGCCACTACGACGCTGAGGTCAAGACCACCTACAAGGCCAAGAAGCCCGT<br>GCAGCTGCCCGGCGCCTACAACGTCAACATCAAGTTGGACATCACCTCCCAC<br>AACGAGGACTACACCATCGTGGAACAGTACGAACGCGCCGAGGGCCGCCAC<br>TCCACCGGCGGCATGGACGAGCTGTACAAGGGCTCCGGCGCCACCAACTTC<br>TCCCTGCTGAAGCAGGCCGGCGACGTGGAGGAGAACCCCGGCCCCCACCT<br>TCAGCAGTCCCCCGTCTACGTGCCGAGCAGCAGGGCCGTACCGCACCAATA<br>TTCCCCGGCGGCAGGCACCCATTTCGGCGCCGCCGCCCACCAGGGAGGCT<br>GGGCTCACGCCGGCGGTTCCTACGGCGACATGGCCTCGCAGGCGCACGGT<br>CTTGGGGGCGCCGCTCACGCCTCTCCCCTCTCCGCCGGTCAGTTCTACACC<br>CAGAACATGGTCATGTCCTCCTGGCGGGCCTACGACGGCTCTGGATTCCAA<br>CGGACGTCACCTTATGGTAAATATCGTCAATATATTTGGATAGTGTATATTAAA<br>TTCAGCATTGGCATGAAGTATGGCGAATCCGAGAAGCCGATTA | mCherry | S | 1322 | 193 | 774 | 355 | 1 | -2 | AS |
| Haxy-h-3-1xHA | T2-1xHA | *Harmonia axyridis* | *h* | CCAAGGATACCCTTACCAGCAAATCTTCGGGTTTCCAGGTGCTGGACCGACA<br>AACCCAGAATTAGCGTATCATCACCAACATCACGTAACAGCTTCTGCCAAGCT<br>TATGGCTACGACATACCCATACGATGTTCCTGACTATGCGTAAACAAGACAAG<br>ACAAGACTGGGAGAATCAACAAAGACA | 1xHA | S | 185 | 118 | 27 | 40 | 0 | 0 | AS |
| Haxy-h-3-3xHA, linker | T2-3xHA | *Harmonia axyridis* | *h* | ACATCACGTAACAGCTTCTGCAAAACTGATGGCTACGACAGGAGGGTCCGGA<br>GGCTACCCATACGATGTTCCTGACTATGCGGGCTATCCCTATGACGTCCCGG<br>ACTATGCAGGATCCTATCCATATGACGTTCCAGATTACGCTTGATGGCTACGA<br>CATAAACAAGACAAGACAAGACTGGGAG | 3xHA | S | 185 | 40 | 105 | 40 | 0 | 0 | AS |
| Haxy-h-3-3xFLAG | T2-3xFLAG | *Harmonia axyridis* | *h* | GTATCATCACCAACATCACGTAACAGCTTCTGCCAAACTGGGCGGCTCCGGC<br>GGCTCCGACTACAAAGACCATGACGGTGATTATAAAGATCATGATATCGATTA<br>CAAGGATGACGATGACAAGATGGCTACGACATAAACAAGACAAGACAAGACT<br>GGGAGAA | 3xFLAG | S | 164 | 40 | 84 | 40 | 2 | -1 | AS |
| P2A, mNeonGreen, linke | T2-mNG | *Harmonia axyridis* | *h* | GCAAATCTTCGGGTTCCCAGGTGCTGGACCGACAAACCCAGAATTAGCGTAT<br>CATCACCAACATCACGTAACAGCTTCTGCAAAACTGATGGCTACGACAGGCT<br>CCGGCGCCACCAACTTCTCCCTGCTGAAGCAGGCCGGCGACGTGGAGGAG<br>AACCCCGGCCCCATGGTGAGCAAGGGCGAGGAGGATAACATGGCCTCTCTC<br>CCAGCGACACATGAGTTACACATCTTTGGCTCCATCAACGGTGTGGACTTTG<br>ACATGGTGGGTCAGGGCACCGGCAATCCAAATGATGGTTATGAGGAGTTAAA<br>CCTGAAGTCCACCAAGGGTGACCTCCAGTTCTCCCCCTGGATTCTGGTCCCT<br>CATATCGGGTATGGCTTCCATCAGTACCTGCCCTACCCTGACGGGATGTCGC<br>CTTTCCAGGCCGCCATGGTAGATGGCTCCGGATACCAAGTCCATCGCACAAT<br>GCAGTTTGAAGATGGTGCCTCCCTTACTGTTAACTACCGCTACACCTACGAG<br>GGAAGCCACATCAAAGGAGAGGCCCAGGTGAAGGGGACTGGTTTCCCTGCT<br>GACGGTCCTGTGATGACCAACTCGCTGACCGCTGCGGACTGGTGCAGGTCG<br>AAGAAGACTTACCCCAACGACAAAACCATCATCAGTACCTTTAAGTGGAGTTA<br>CACCACTGGAAATGGCAAGCGCTACCGGAGCACTGCGCGGACCACCTACAC<br>CTTTGCCAAGCCAATGGCGGCTAACTATCTGAAGAACCAGCCGATGTACGTG<br>TTCCGTAAGACGGAGCTCAAGCACTCCAAGACCGAGCTCAACTTCAAGGAGT<br>GGCAAAAGGCCTTTACCGATGTGATGGGCATGGACGAGCTGTACAAGGGCG<br>GCAGCGGCGGCAGCCCGAAGAAGAAACGAAAGGTTGACCCTAAGAAAAAGA<br>GGAAGGTGGACCCCAAGAAAAAGCGAAAATAAACAAGACAAGACAAGACTGG<br>GAGAATCAACAAAGACACTAACCTTGCTGCGCTATAAAATGAGGAGAGTTTTA<br>TTTGTAGTAGCAGCAGATTCCGAT | mNeonGreen | S | 1058 | 166 | 726 | 166 | 0 | -14 | AS |
| Haxy_h-intron-attP | T3-attP | *Harmonia axyridis* | *h* | TCCTGCGAAAGAATCGAAAAAACACAATGCAGAGTTATATTTTTTTTTACAAAA<br>AGCAATCGTGCAGGTGCGTGCCCCAACTGGGGTAACCTTTGAGTTCTCTCAG<br>TTGGGGGAGTTGGTGTTTAGGGATTCACAGCTTGGCTTGATTT | attP | AS | 149 | 36 | 42 | 71 | 0 | 0 | AS |
| Haxy-h-3ext | T2-3xFLAG-3´ext-ds | *Harmonia axyridis* | *h* | TGATGGCTACGACAGGAGGGTCCGGAGGCGACTACAAAGACCATGACGGTG<br>ATTATAAAGATCATGATATCGATTACAAGGATGACGATGACAAGTGATGGCTA<br>CGACATAAACAAGACAAGACAAGACTGGGAG | 3xFLAG | S | 135 | - | 95 | 40 | 0 | 0 | AS |
| Haxy-h-3ext(-) | | *Harmonia axyridis* | *h* | CTTGTCATCGTCATCCTTGTAATCGATATCATGATCTTTATAATCACCGTCATG<br>GTCTTTGTAGTCGCCTCCGGACCCTCCTGTCGTAGCCATCAGTTTTGCAGAA<br>GCTGTTACGTGATGTTGGTGATGATACGC | 3xFLAG | AS | 135 | 40 | 95 | - | | | AS |
| Haxy-h-blunt | T2-3xFLAG-blunt-ds | *Harmonia axyridis* | *h* | ACATCACGTAACAGCTTCTGCAAAACTGATGGCTACGACAGGAGGGTCCGGA<br>GGCGACTACAAAGACCATGACGGTGATTATAAAGATCATGATATCGATTACAA<br>GGATGACGATGACAAGTGATGGCTACGACATAAACAAGACAAGACAAGACTG<br>GGAG | 3xFLAG | S | 161 | 40 | 81 | 40 | 14 | 0 | AS |
| Haxy-h-blunt | | *Harmonia axyridis* | *h* | CTCCCAGTCTTGTCTTGTCTTGTTTATGTCGTAGCCATCACTTGTCATCGTCA<br>TCCTTGTAATCGATATCATGATCTTTATAATCACCGTCATGGTCTTTGTAGTCG<br>CCTCCGGACCCTCCTGTCGTAGCCATCAGTTTTGCAGAAGCTGTTACGTGAT<br>GT | 3xFLAG | AS | 161 | 40 | 81 | 40 | | | AS |
| Haxy-h-5ext | T2-3xFLAG-5´ext-ds | *Harmonia axyridis* | *h* | GCGTATCATCACCAACATCACGTAACAGCTTCTGCAAAACTGATGGCTACGA<br>CAGGAGGGTCCGGAGGCGACTACAAAGACCATGACGGTGATTATAAAGATC<br>ATGATATCGATTACAAGGATGACGATGACAAG | 3xFLAG | S | 135 | 40 | 95 | - | 0 | 0 | AS |
| Haxy-h-5ext(-) | | *Harmonia axyridis* | *h* | CTCCCAGTCTTGTCTTGTCTTGTTTATGTCGTAGCCATCACTTGTCATCGTCA<br>TCCTTGTAATCGATATCATGATCTTTATAATCACCGTCATGGTCTTTGTAGTCG<br>CCTCCGGACCCTCCTGTCGTAGCCATCA | 3xFLAG | AS | 135 | - | 95 | 40 | | | AS |
| Haxy-en | T4-3xFLAG-3´ext-ds | *Harmonia axyridis* | *engrailed* | GGAGGGTCCGGAGGCGACTACAAAGACCATGACGGTGATTATAAAGATCAT<br>GATATCGATTACAAGGATGACGATGACAAGTAGACATCGAGGAGTACTGCAA<br>GTCTATTTGATGAATAAACCCAGTGA | 3xFLAG | S | 129 | - | 89 | 40 | -3 | 5 | AS |
| Haxy-en(-) | | *Harmonia axyridis* | *engrailed* | GATGTCTACTTGTCATCGTCATCCTTGTAATCGATATCATGATCTTTATAATCA<br>CCGTCATGGTCTTTGTAGTCGCCTCCGGACCCTCCGCTCATAGGCATGTCAT<br>CTTCGTCGCAAGGTACTGTGGAG | 3xFLAG | AS | 129 | 40 | 89 | - | | | AS |
| Gbim-piwi | T5-3xFLAG-3´ext-ds | *Gryllus bimaculatus* | *piwi* | GTATTATTTGGGAGGGTCCGGAGGCGACTACAAAGACCATGACGGTGATTAT<br>AAAGATCATGATATCGATTACAAGGATGACGATGACAAGTAAGTTGTGTTACA<br>TTCCTATATGTGAAGACTTTTTTAATATTCACTTTACTTC | 3xFLAG | S | 145 | - | 105 | 40 | top:-20/bottom:-25 | top:-3/bottom:+2 | AS |
| Gbim-piwi(-) | | *Gryllus bimaculatus* | *piwi* | TGTAACACAACTTACTTGTCATCGTCATCCTTGTAATCGATATCATGATCTTTA<br>TAATCACCGTCATGGTCTTTGTAGTCGCCTCCGGACCCTCCCAAATAATACA<br>GGTAATTCTCTAATTGAGTATTTGGAGCCCTGTGCAAAC | 3xFLAG | AS | 145 | 40 | 105 | - | | | AS |

Table S3. Primers used for mutation screening

| primer name | Sequence |
|---|---|
| Haxy_en_3'_Fw | CAGCTCATGGCCCAAGGG |
| Haxy_en_3'_Rv | CAGTATGAGATCTTTCACTGCACG |
| Gbim_piwi_3'_Fw | TGGCTTACTTAGTTGGACAG |
| Gbim_piwi_3'_Rv | CTCAGAACATCCATCATTGAAC |
| Haxy_h_3'_Fw | ATACGATTACAGCTGCCTGC |
| Haxy_h_3'_Rv | AGAGTATCGGAATCTGCTGC |
| Haxy_h_5'_Fw | ATGTTCCACACCGGCG |
| Haxy_h_5'_Rv | TTCTGGGTGTAGAACTGACC |
| Haxy_h_amplicon_F | AACCAAGGATACCCTTACC |
| Haxy_h_amplicon_R | AGTGTCTTTGTTGATTCTCC |

Table S4. p-values of chi-square tests of the data in Table S6

| | | blunt-fast | blunt-slow | 5'ex-fast | 5'ex-slow | 3'ext-fast | 3'ext-slow | no anneal | ssDNA-S |
|---|---|---|---|---|---|---|---|---|---|
| NHEJ | blunt-slow | 0 | - | - | - | - | - | - | - |
| | 5'ex-fast | 0 | 1.20E-270 | - | - | - | - | - | - |
| | 5'ex-slow | 0 | 5.19555480641 | 0.010215 | - | - | - | - | - |
| | 3'ext-fast | 0 | 0 | 0 | 0 | - | - | - | - |
| | 3'ext-slow | 0 | 0 | 0 | 1.85E-299 | 0 | - | - | - |
| | no anneal | 0 | 0 | 0 | 0 | 1.56E-161 | 0 | - | - |
| | ssDNA-S | 0 | 0 | 0 | 0 | 0 | 0 | 0 | - |
| | ssDNA-AS | 0 | 3.61E-188 | 1.40E-08 | 1.21E-16 | 0 | 0 | 0 | 0 |
| Precise | blunt-slow | 5.26483E-28 | - | - | - | - | - | - | - |
| | 5'ex-fast-fast | 2.44461E-56 | 2.66E-11 | - | - | - | - | - | - |
| | 5'ex-fast-slow | 4.45811E-45 | 4.47392E-05 | 0.081405 | - | - | - | - | - |
| | 3'ext-fast | 5.85409E-68 | 8.78106E-22 | 0.000673 | 6.08875E-09 | - | - | - | - |
| | 3'ext-slow | 1.45007E-30 | 0.110798619 | 0.000103 | 2.47E-01 | 7.56676E-14 | - | - | - |
| | no anneal | 7.18002E-52 | 2.88557E-13 | 0.556591 | 0.001321469 | 1.11E-01 | 5.80258E-07 | - | - |
| | ssDNA-S | 3.99204E-89 | 3.14981E-29 | 1.11E-05 | 2.01052E-12 | 0.824832906 | 7.22835E-19 | 0.013862 | - |
| | ssDNA-AS | 1.41948E-39 | 4.71E-03 | 2.13E-03 | 7.94E-01 | 9.89503E-12 | 0.824832906 | 1.8E-05 | 4.57E-16 |
| Precise + Imprecise | blunt-slow | 3.9841E-09 | - | - | - | - | - | - | - |
| | 5'ex-fast-fast | 9.4357E-209 | 1.56E-290 | - | - | - | - | - | - |
| | 5'ex-fast-slow | 3.50602E-09 | 2.30466E-35 | 3.8E-140 | - | - | - | - | - |
| | 3'ext-fast | 1.1089E-135 | 1.7118E-204 | 5.17E-09 | 7.03073E-81 | - | - | - | - |
| | 3'ext-slow | 1.73522E-07 | 1.84605E-29 | 6.1E-139 | 1.00E+00 | 3.10603E-80 | - | - | - |
| | no anneal | 2.8978E-191 | 9.0936E-259 | 0.240871 | 7.1764E-133 | 7.31E-13 | 1.2852E-132 | - | - |
| | ssDNA-S | 2.39503421073 | 0 | 1.2E-07 | 6.9242E-225 | 1.22681E-32 | 5.0416E-223 | 0.002114 | - |
| | ssDNA-AS | 2.1143E-275 | 0.00E+00 | 1.29E-07 | 1.45E-198 | 1.69432E-30 | 9.6151E-198 | 0.001941 | 1 |

Red, p-value < 0.01, NHEJ, read data mapped to the reference without the insert; Insert, read data mapped to the refference with the insert.
Multiple comparisons were corrected with Holm's method.

Table S5. Statistics of the gene knock-in using ssDNA templates

| DNA template ID | Insertion tag | Injected eggs *n* | Screened adults *n* | Insertion founder pair *n* (%) | Precise insertion *n* | 5´ Fidelity % (*n/n*) | 3´ Fidelity % (*n/n*) |
|---|---|---|---|---|---|---|---|
| T1-1xHA | HA tag, 1x | 579 | 62 | 5 (8.06%) | 4 | 80% (4/5) | 100% (5/5) |
| T1-3xHA | HA tag, 3x | 98 | 9 | 1 (11.11%) | 0 | 0% (0/1) | 100% (1/1) |
| T1-mCh | mCherry-2A | 203 | 33 | 1 (3.03%) | 0 | N.D. | N.D. |
| T2-1xHA | HA tag, 1x | 93 | 14 | 0 (0%) | 0 | - | - |
| T2-3xHA | HA tag, 3x, linker | 255 | 19 | 2 (10.53%) | 0 | 0% (0/5) | 100% (2/2) |
| T2-3xFLAG | FLAG tag, 3x, linker | 277 | 41 | 4 (9.76%) | 0 | 0% (0/5) | 100% (4/5) |
| T2-mNG | P2A, mNeonGreen, linker, 3xNLS | 225 | 40 | 1 (2.5%) | 0 | 0% (0/1) | 0%** (0/1) |
| T3-attP | attP* | 45 | 19 | 2 (10.53%) | 1 | 50% (1/2) | 100% (2/2) |

*: injected to *h/hC* heterozygote eggs to target only a single *pannier* allele (*h*)

**:a partial fragment was inserted in the reverse direction

N.D: not determined

Table S6. Read counts of the deep sequencing analysis

| Injected DNA template | Reference with insert | | | Reference without insert | | Total aligned reads |
|---|---|---|---|---|---|---|
| | Precise | Precise+Imprecise | Other | NHEJ | Other | |
| blunt-fast | 381 (0.52%) | 2044 (2.79%) | 71125 (97.2%) | 54080 (73.9%) | 19089 (26.1%) | 73169 |
| blunt-slow | 165 (0.19%) | 2901 (3.34%) | 84014 (96.7%) | 29468 (33.9%) | 57447 (66.1%) | 86915 |
| 5'ex-fast | 40 (0.06%) | 445 (0.64%) | 68805 (99.4%) | 29499 (42.6%) | 39751 (57.4%) | 69250 |
| 5'ex-slow | 69 (0.1%) | 1607 (2.27%) | 69162 (97.7%) | 30628 (43.3%) | 40141 (56.7%) | 70769 |
| 3'ext-fast | 9 (0.01%) | 610 (0.94%) | 64024 (99.1%) | 14524 (22.5%) | 50110 (77.5%) | 64634 |
| 3'ext-slow | 81 (0.14%) | 1373 (2.31%) | 58143 (97.7%) | 31884 (53.6%) | 27632 (46.4%) | 59516 |
| 3'ext-no anneal | 21 (0.04%) | 315 (0.56%) | 55565 (99.4%) | 9080 (16.2%) | 46800 (83.8%) | 55880 |
| ssDNA-S | 9 (0.01%) | 361 (0.43%) | 83428 (99.6%) | 4532 (5.4%) | 79257 (94.6%) | 83789 |
| ssDNA-AS | 84 (0.12%) | 300 (0.42%) | 70668 (99.6%) | 29147 (41.1%) | 41821 (58.9%) | 70968 |
